# A generalist lifestyle allows rare *Gardnerella* spp. to persist at low levels in the vaginal microbiome

**DOI:** 10.1101/2020.08.25.267146

**Authors:** Salahuddin Khan, Sarah J. Vancuren, Janet E. Hill

## Abstract

*Gardnerella* spp. are considered a hallmark of bacterial vaginosis, a dysbiosis of the vaginal microbiome. There are four cpn60 sequence-based subgroups within the genus (A, B, C, and D), and thirteen genome species have been defined recently. *Gardnerella* spp. co-occur in the vaginal microbiome with varying abundance, and these patterns are shaped by a resource-dependent, exploitative competition, which affects the growth rate of subgroup A, B, and C negatively. The growth rate of rarely abundant subgroup D, however, increases with the increasing number of competitors, negatively affecting the growth rate of others. We hypothesized that a nutritional generalist lifestyle and minimal niche overlap with the other, more abundant *Gardnerella* spp. facilitate the maintenance of subgroup D in the vaginal microbiome through negative-frequency dependent selection. Using 40 whole genome sequences from isolates representing all four subgroups we found that they could be distinguished based on content of their predicted proteomes. Proteins associated with carbohydrate and amino acid uptake and metabolism were significant contributors to the separation of subgroups. Subgroup D isolates had significantly more of their proteins assigned to amino acid metabolism than the other subgroups. Subgroup D isolates were also significantly different from others in terms of number and type of carbon sources utilized in a phenotypic assay, while the other three could not be distinguished. Overall, the results suggest that a generalist lifestyle and lack of niche overlap with other *Gardnerella* spp. leads to subgroup D being favoured by negative-frequency dependent selection in the vaginal microbiome.

## Introduction

*Gardnerella* spp. are an important diagnostic marker of bacterial vaginosis (BV), a dysbiosis of the vaginal microbiome characterized by a shift from lactobacilli dominated vaginal microbiome to a more diverse microbiome, containing many aerobic and anaerobic bacterial species, including *Gardnerella* spp‥ *Gardnerella* is a diverse genus and at least four subgroups (A, B, C, and D) have been identified using cpn60 universal target barcode sequencing [1], which correspond to four clades defined by Ahmed et al. [2]. Recently, *Gardnerella* subgroups have been reclassified into thirteen genome species, of which four are now named as *G. vaginalis* (Subgroup C/Clade 1), *G. swidsinskii* and *G. leopoldii* (Subgroup A/Clade 4), and *G. piotii* (Subgroup B/Clade 2) [3, 4]. These *Gardnerella* species differ in their phenotypic traits, including sialidase activity and vaginolysin production, which may render some of the subgroups more pathogenic than the others [5–7].

Women with vaginal microbiomes dominated by *Gardnerella* are usually colonized by at least two *Gardnerella* spp. [4, 8]. The relative abundances of these co-occurring species, however, are not equal. Subgroup A (*G. swidsinksii* and *G. leopoldii*) and subgroup C (*G. vaginalis*) are most frequently dominant in reproductive aged women [4, 8]. These two subgroups are also often associated with the clinical symptoms of bacterial vaginosis [4, 9, 10]. Subgroup B has been suggested to be associated with intermediate microbiota [7, 9, 10]. Subgroup D, comprised of several unnamed "genome species", has only been detected at low prevalence and abundance [4, 10].

Several factors can affect the abundance and co-occurrence of *Gardnerella* spp. in the vaginal microbiome, including host physiology, host-microbiota interactions, nutrient availability and ecological interactions among bacteria [11, 12]. Ecological interactions are perhaps the most important factors which may affect the co-occurrence and ecological succession of *Gardnerella* species in the vaginal microbiome. Recently, we demonstrated that an indirect, exploitative competition between subgroups of *Gardnerella* is prevalent in co-cultures *in vitro*. While the growth rates of isolates in subgroups A, B, and C, were negatively affected by competition, growth rates of *Gardnerella* subgroup D isolates increased with the increasing number of competing subgroups in co-culture communities [12].

The strength of microbial interactions between bacterial species can be affected by niche overlap [13, 14], and species with similar nutritional requirements will naturally compete over the same resources [15]. In addition to competition for nutritional resources, bacteria may also compete for resources essential for colonizing a specific site. Since isolates from *Gardnerella* subgroups A, B and C are negatively affected by competition, and subgroup D isolates experienced a boost in growth rate, the degree of niche overlap between subgroup A, B, and C is presumably higher than between subgroup D and any of the others.

Although the growth rate of subgroup D increases in co-cultures, it does not have an intrinsically high growth rate. In fact, the *in vitro* growth rate of subgroup D is half of that of subgroup C, which may contribute to its low abundance in the vaginal microbiome [12]. Low abundance species are often favoured by negative-frequency-dependent selection [16, 17], which can be governed by nutritional requirements [18]. Bacteria capable of utilizing relatively few, abundantly available nutrients in a particular environment are nutritional specialists in the context of that environment. Generalists, on the contrary, are bacteria capable of utilizing more nutrient sources than their specialist counterparts. In negative frequency-dependent selection, the resources accessible to rapidly growing specialists will dwindle, reducing the fitness of the specialists as their population increases. As a result, the population of more generalist bacteria capable of utilizing a wider range of nutrient sources will expand in a density-dependent manner [18, 19]. Generalists can also negatively affect the growth of specialists by competing for the resources that can be utilized by both of them [14].

Although growth of the rarely abundant subgroup D is facilitated in co-cultures, the degree of overlap in nutrient utilization among the subgroups and the range of nutrient utilization by individual subgroups are yet unknown. The objective of our present study was, therefore, to evaluate the amount of genomic and phenotypic overlap in nutrient utilization among the subgroups of *Gardnerella* and to determine if subgroup D is a nutritional generalist relative to the three other subgroups. Findings are interpreted in relation to the hypothesis that subgroup D is maintained in the vaginal microbiome through negative frequency dependent selection.

## Methods

### Bacterial isolates

Thirty-nine *Gardnerella* isolates from our culture collection representing all four subgroups (based on cpn60 barcode sequencing) were selected for the study (*n* = 12 subgroup A, 12 subgroup B, 8 subgroup C, and 7 subgroup D isolates) (Table S1). Isolates were streaked on Columbia agar plates with 5% (v/v) sheep blood and were incubated anaerobically at 37° C for 48 h. For broth culture, colonies from blood agar plates were suspended in BHI broth supplemented with 10% horse serum and 0.25% (w/v) maltose.

### Whole-genome sequencing

Whole genome sequences for 10 of the study isolates had been published previously, and the remaining 29 were sequenced as part of the current study (Table S1). DNA was extracted from isolates using a modified salting out protocol [20] and was stored at −20°C. DNA was quantified using Qubit dsDNA BR assay kit (Invitrogen, Burlington, Ontario) and the quality of the extracts was assessed by the A260/A280 ratio. Isolate identity was confirmed by cpn60 barcode sequencing as follows. cpn60 barcode sequences were amplified from extracted DNA with the primers JH0665 (CGC CAG GGT TTT CCC AGT CAC GAC GAY GTT GCA GGY GAY GGH CHA CAA C) and JH0667 (AGC GGA TAA CAA TTT CAC ACA GGA GGR CGATCR CCR AAK CCT GGA GCY TT). The reaction contained 2 μL template DNA in 1× PCR buffer (0.2 M Tris-HCl at pH 8.4, 0.5 M KCl), 2.5 mM MgCl2, 200 μM dNTP mixture, 400 nM of each primer, 2 U AccuStart Taq DNA polymerase, and water to bring to a final volume of 50 μL. PCR was carried out with incubation at 94°C for 30 seconds, 40 cycles of 94°C 30 sec, 60°C for 1 min, 72°C for 1min, and final extension at 72°C for 10 min. PCR products were purified and sequenced by Sanger sequencing and compared with the chaperonin sequence database cpnDB [21] to confirm identity.

Following confirmation of the identity of isolates, sequencing libraries were prepared using the Nextera XT DNA library preparation kit according to the manufacturer’s instructions (Illumina, Inc., San Diego, CA). PhiX DNA (15% [vol/vol]) was added to the indexed libraries before loading onto the flow cell. The 500 cycle V2 reagent kit was used for the Illumina MiSeq platform (Illumina, Inc.).

Raw sequences were trimmed using Trimmomatic [22] with a minimum quality score of 20 over a sliding window of 4, and minimum read length of 40. Trimmed sequences were assembled using SOAPdenovo2 [23] or SPAdes (NR002, NR043, NR044) [24]. Assembled genomes were annotated using the National Center for Biotechnology Information Prokaryotic Genome Annotation Pipeline [25].

### Pangenome analysis

Pangenome analysis of the 39 study isolates and the published genome of *G. vaginalis* strain ATCC 14019 (Accession number: PRJNA55487) was performed using the micropan R package [26]. We used “complete” linkage for clustering, and the cut-off value for the generation of clusters was set to 0.75. For initial visualization of the results, the Jaccard index was used to calculate similarity of patterns of presence and absence of protein clusters among all isolates and a dendrogram was constructed from the results by unweighted pair group method with arithmetic mean (UPGMA) using DendroUPGMA (http://genomes.urv.cat/UPGMA/).

### COG analysis

Predicted protein sequences from individual genomes were classified into Clusters of Orthologous Groups (COG) categories using WebMEGA (http://weizhonglab.ucsd.edu/webMEGA). Based on the output from this process, the proportion of proteins in each of the COG categories was calculated for each genome. The distributions of proportional abundances of each category were then used to assess the relationships of the four subgroups in terms of COG category representation.

### Carbon source utilization assay

Bacterial isolates from freezer stocks were streaked on 5% sheep blood agar plates and were grown for 48 h anaerobically, prior to inoculation of AN Microplates (Biolog Inc, Hayward, CA). Each plate contained 95 carbon sources and one blank well. Colonies of *Gardnerella* isolates were harvested using a sterile swab and suspended in 14 mL of inoculating fluid supplied by the manufacturer. The cell density was adjusted to 55%T (OD_595_ approximately 0.25) using a turbidimeter. Each well was filled with 100 μl of culture suspension and was incubated at 35° C anaerobically for 48 h. All inoculations and incubations were performed in an anaerobic chamber containing 10 % CO_2_, 5% hydrogen, and 85% nitrogen. All plates were read visually after 48h of incubation. If there was no carbon source utilization, the wells remained colourless. A visual change from colourless to purple indicated carbon source utilization. To avoid bias in interpretation, a subset of the plates was read by a second observer who was blinded to the identity of the isolates. There was no disagreement between independent observers. The entire experiment was performed in two biological replicates.

### Carbon source profiling of co-cultures

Representative isolates (VN003 of subgroup A, VN002 of subgroup B, NR001 of subgroup C, and WP012 of subgroup D) from the four subgroups were co-cultured in the Biolog AN Microplate in a pairwise fashion (n =6, AB, AC, AD, BC, BD, CD), by combining 50 μL of each isolate suspended in inoculation fluid in each well. The co-cultured AN Microplates were incubated at 35°C for 48h before being assessed visually for colour change. The experiment was repeated on separate days.

### Statistical analysis

The degree of similarity between the isolates in terms of presence/absence of protein clusters generated in the pangenome analysis, proportional abundance of proteins in various COG categories, and carbon source utilization patterns were calculated using the Bray-Curtis dissimilarity matrix. Principle components analysis (PCA) was performed on the distance matrices and significance of relationships were tested using PERMANOVA with the ADONIS function in the *vegan* package [27]. The SIMPER function was used to identify variables driving the differences between groups.

One-way ANOVA, student’s t-test and chi-square tests were applied to determine if utilization of particular carbon sources was associated with specific subgroups.

All statistical analyses were performed in RStudio (version 3.5.2). Figures were generated using GraphPad Prism 8.0 and RStudio (version 3.5.2).

## Results

### Overlap between the subgroups based on pangenome and COG analysis

The purpose of our pangenome analysis was to estimate the degree of niche overlap between *Gardnerella* subgroups based on comparisons of their predicted proteomes. Hierarchical clustering using complete linkage produced 4,868 clusters or predicted proteins in the pangenome of the 40 isolates included (rarefaction curve shown in Fig. S1). The strict core (defined as the protein clusters present in all isolates) included 176 clusters. Most of these core proteins were related to metabolism, transcriptional control, DNA replication, and protein synthesis. Clustering of the genomes by subgroup was apparent in a UPGMA dendrogram based on the presence/absence patterns of the 4,868 protein clusters (Fig. 1a). PCA was performed to determine the extent of overlap between the subgroups. The amount of variance explained by the two principal components was 19.4%, based on which, the four subgroups were separable (Fig. 1b). The dissimilarity between the four subgroups was significant (pairwise-ADONIS, Bonferroni adjusted, p < 0.05, A vs B, R^2^ = 0.45; A vs C, R^2^ = 0.48; A vs D, R^2^ = 0.26; B vs C, R^2^ = 0.34; B vs D, R^2^ = 0.45; and C vs D, R^2^ = 0.55).

**Fig. 1.**
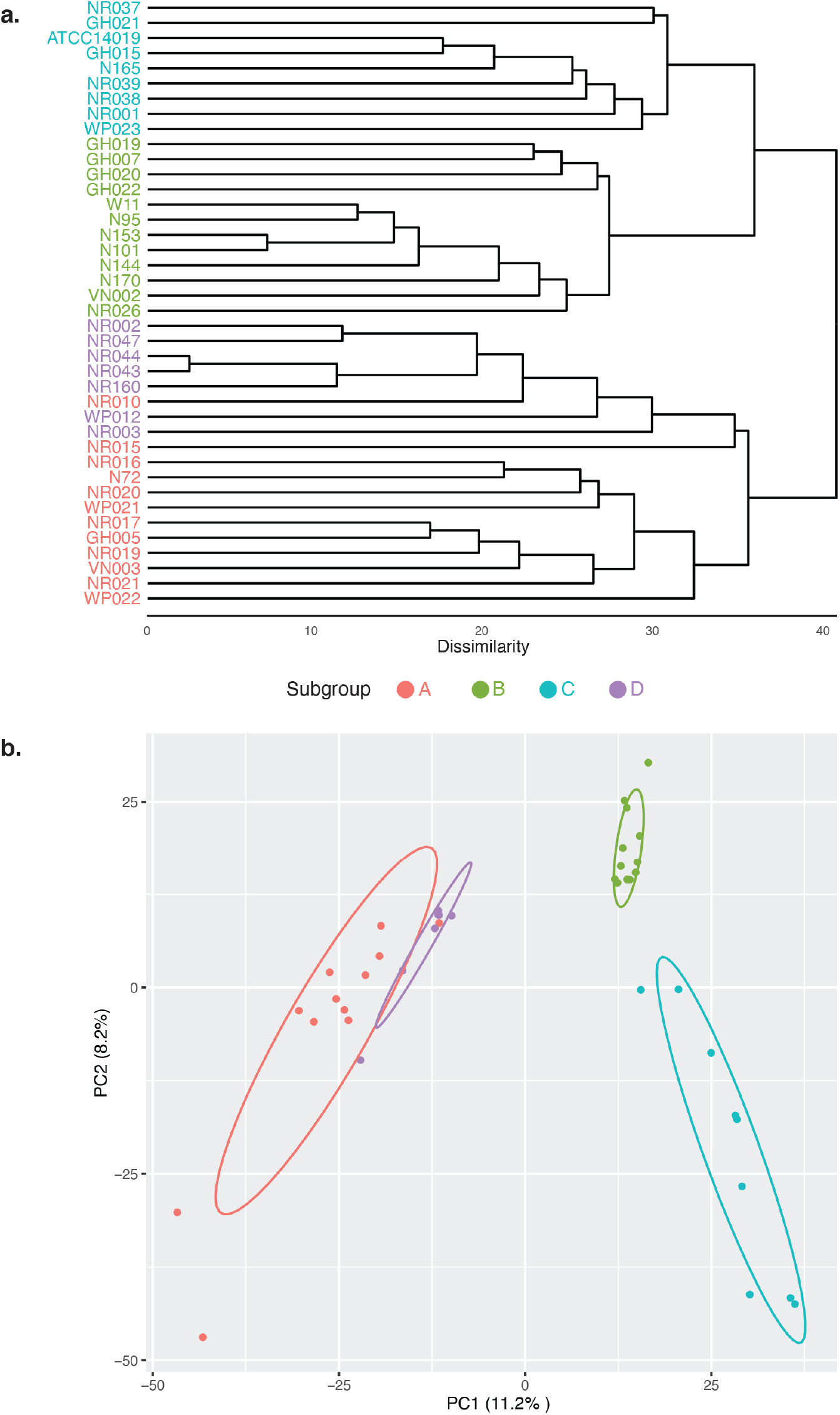
Comparison of predicted proteomes of study isolates. **(a)** UPGMA dendrogram based on presence/absence of protein clusters in the predicted proteomes of *Gardnerella* isolates. **(b)** Principle components analysis (PCA) of Bray-Curtis dissimilarity matrices calculated from protein cluster distributions. Dissimilarity between the four subgroups is significant (pairwise ADONIS p < 0.05, Bonferroni adjusted). Subgroup affiliations of isolates are indicated by colour as shown in the legend between the panels.

Following the identification of core and accessory proteins, we investigated the distribution of functional classifications of proteins encoded by isolates in the four subgroups. COG analysis resulted in assignment of predicted proteins into 23 functional categories. As expected, hierarchical clustering of the COG distribution patterns corresponded to subgroup affiliation (Fig. 2a). PCA was performed on the Bray-Curtis dissimilarity matrix and the differences between all subgroups were found to be significant (pairwise ADONIS, Bonferroni adjusted, p <0.05, A vs B, R^2^ = 0.31; A vs C, R^2^ = 0.71; A vs D, R^2^ = 0.26; B vs C, R^2^ = 0.48; B vs D, R^2^ = 0.20; and C vs D, R^2^ = 0.74) (Fig. 2b).

**Fig. 2.**
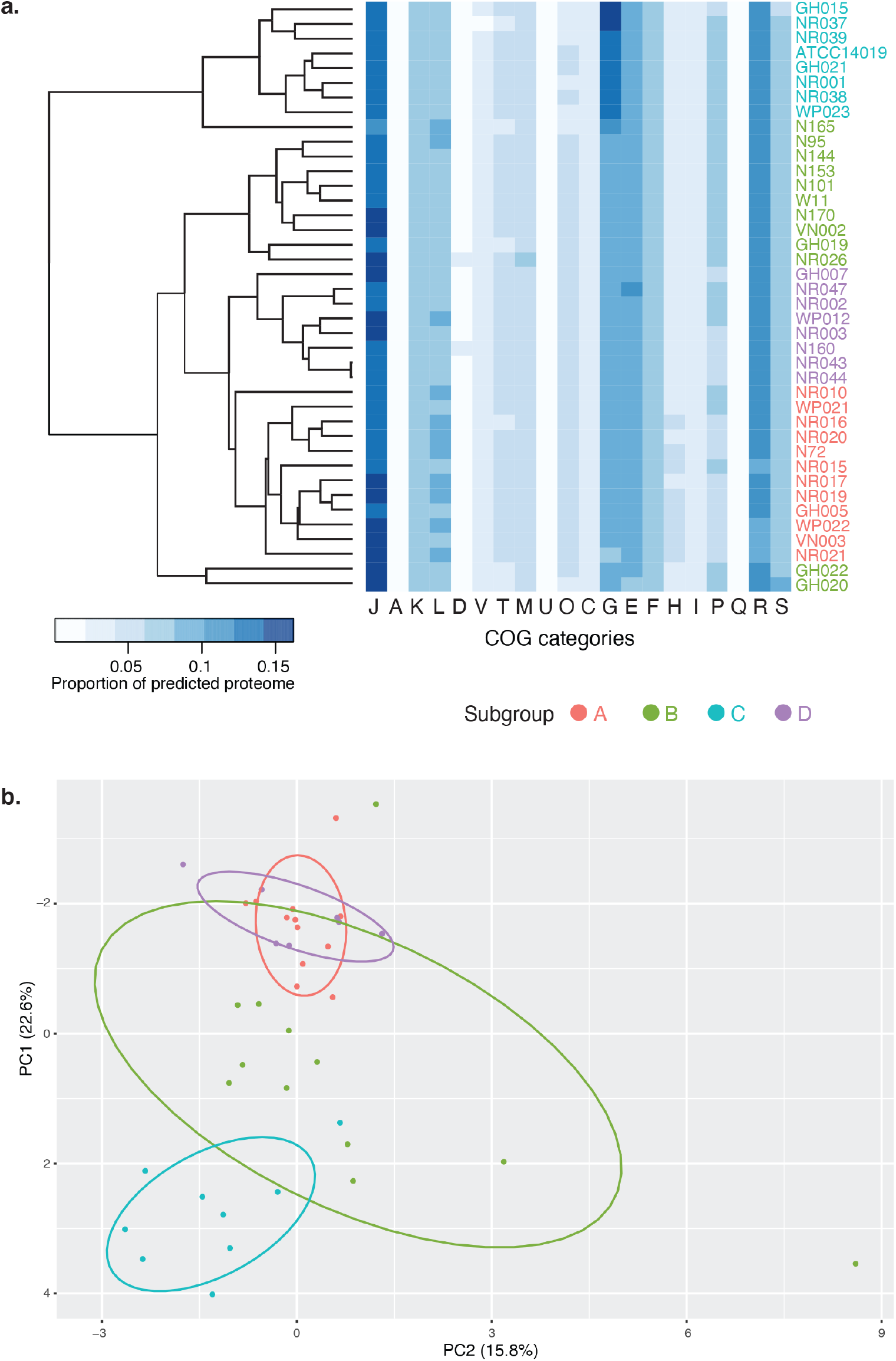
COG analysis of predicted proteomes. **(a)** Hierarchical clustering of study isolates based on proportional abundance of COG categories in their predicted proteomes. Abundance is indicated by blue colour intensity in the heat map. **(b)** PCA of Bray-Curtis dissimilarity matrices calculated from the proportional abundance data. Dissimilarity between the four subgroups is significant (pairwise ADONIS, p < 0.05, Bonferroni adjusted). Subgroup affiliations of isolates are indicated by colour as shown in the legend between the panels.

We identified the variables which caused the four subgroups to diverge in terms of abundance of different COG categories in a multivariate analysis using SIMPER [28]. SIMPER calculates the contribution of each variable to the dissimilarity observed between two groups and relies on Bray-Curtis dissimilarity matrix for calculating the proportion of contribution of each variable being tested. Thirty-six percent of the differences between subgroup A and subgroup B were accounted for by amino acid transport and metabolism (COG category E), inorganic ion transport metabolism (category P), translation, ribosomal structure and biogenesis proteins (category J). The proportion of proteins with functions related to carbohydrate transport and metabolism (category G) was the major factor that differentiated subgroup A from C, contributing to 34% of the dissimilarity observed. Carbohydrate transport and metabolism also accounted for 31% of the dissimilarity observed between subgroups B and C and 36% of the dissimilarity between subgroups C and D. The major contributing factors that differentiated subgroup A and D were proportional abundance of proteins assigned to functional categories H (co-enzyme transport and metabolism) and E (amino acids transport and metabolism) (23%). Subgroups B from D were separated primarily based on functional categories P (inorganic ion transport and metabolism), J (translation, ribosomal structure and biogenesis), G (carbohydrate transport and metabolism proteins), and E (amino acid transport and metabolism proteins), which together accounted for 37% of the dissimilarity observed.

### Functional categories of proteins differentiating subgroups of *Gardnerella*

We tested if the proportions of individual functional categories of proteins that drive the overall separation of the four subgroups in multivariate analysis were significantly different between pairs of *Gardnerella* subgroups. This analysis revealed that subgroup C has a significantly higher proportion of its encoded proteins associated with carbohydrate transport and metabolism and transport, and transcriptional regulation than the other subgroups (unpaired t-test, p ≤ 0.01, Bonferroni adjusted, Fig. 3a, 3d). The proportion of proteins associated with amino acid transport and metabolism is significantly higher in subgroup D than subgroups A, B, and C (unpaired t-test, p ≤ 0.01, Bonferroni adjusted, Fig. 3b). Proteins involved in co-enzyme transport and metabolism were found in significantly higher proportional abundance in subgroup A than in subgroup B, C and D (unpaired t-test, p ≤ 0.0001, Bonferroni adjusted, Fig. 3c). Subgroup B has a significantly higher abundance of proteins associated with inorganic ion transport and metabolism than subgroup A and D (unpaired t-test, p ≤ 0.0001, Fig. 3e), but the difference between subgroup B and C was not significant. Subgroup B also has a significantly higher proportion of translation, ribosomal structure and biogenesis proteins (unpaired t-test, p ≤ 0.001, Fig. 3f) compared to subgroup C.

**Fig. 3.**
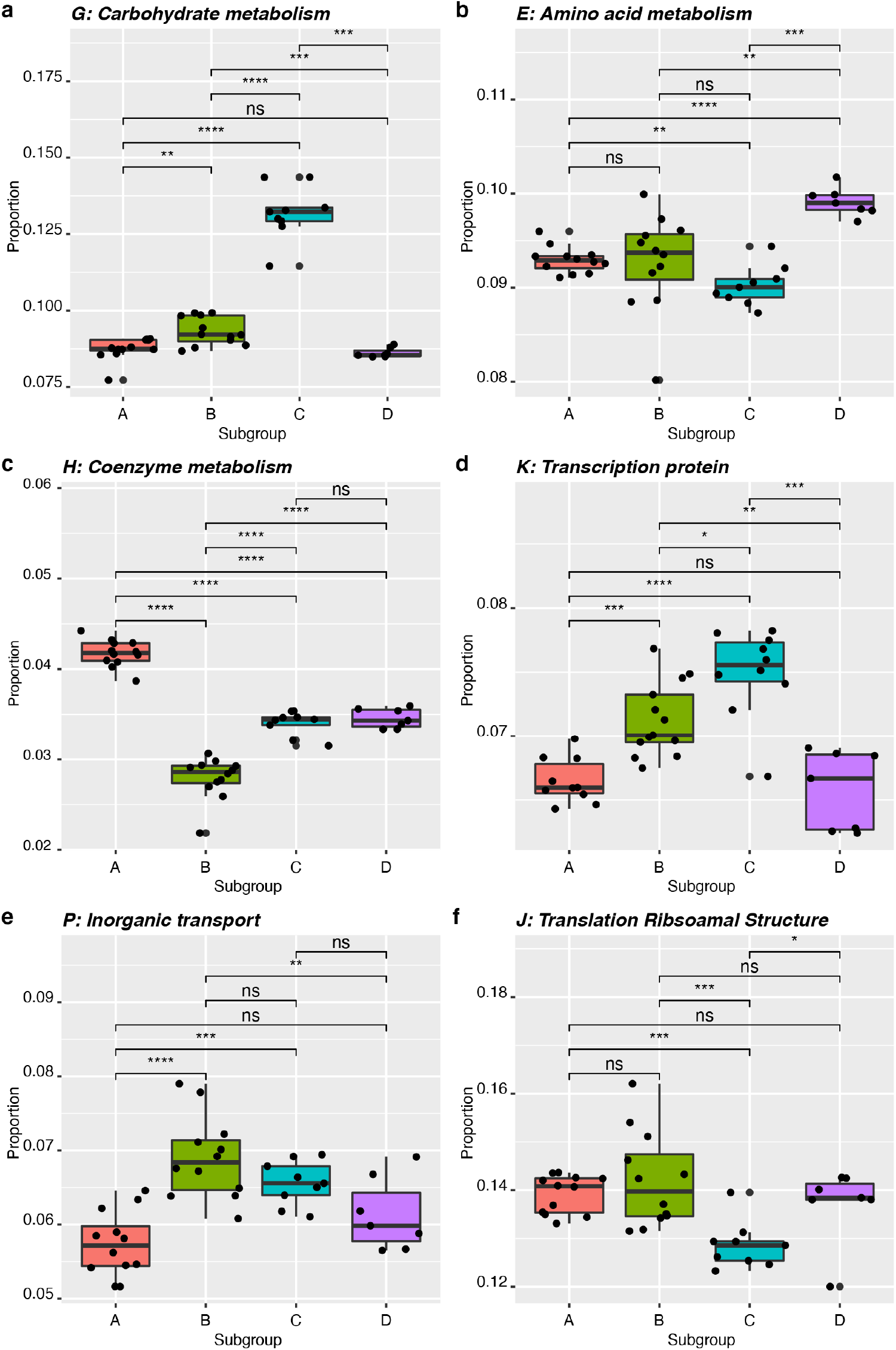
Differential abundance of six COG categories that were identified by SIMPER analysis as main drivers of subgroup separation. **(a)** carbohydrate metabolism and transport proteins (Category G), **(b)** amino acid transport and metabolism proteins (Category E), **(c)** co-enzyme transport and metabolism proteins (Category H), **(d)** transcription proteins (Category K), **(e)** inorganic ion transport and metabolism proteins (Category P), and **(f)** translation, ribosomal structure and biogenesis proteins (Category J). Results of unpaired t-tests are indicated where * is p ≤ 0.05, ** is p ≤ 0.01, *** is p ≤ 0.001, **** is p ≤ 0.0001 and ns is not significant.

### Carbon source utilization phenotypes

We hypothesized that subgroup D, a slow-growing, rarely detected *Gardnerella* subgroup is maintained in the vaginal microbiome at a low level and avoids competitive exclusion through negative-frequency-dependent selection, made possible by being a nutritional generalist. We performed carbon source utilization profiling of thirty-six representative isolates (n= 12, subgroup A; n= 9, subgroup B; n=8, subgroup C (including type strain *G. vaginalis* ATCC 14018); and n=7, subgroup D). The number of carbon sources utilized by any *Gardnerella* strain ranged from 5 to 24. Only 25% (9/36) of the isolates utilized more than 17 carbon sources, including two subgroup C (NR001, NR038) and all subgroup D isolates. Twenty isolates utilized at least 13 carbon sources, including three subgroup A (3/12, 25%), four subgroup B (4/8, 50%), six subgroup C (6/7, 86%), and all seven isolates of subgroup D (100%). The average number of carbon sources utilized by isolates in subgroups A, B, C, and D was 10.4±3.1, 11.8±1.75, 13.9±3.6, and 20.3±1.9, respectively (Fig. 4). A one-way ANOVA was performed to compare the overall difference in carbon sources utilization among the four subgroups showed significant difference among the subgroups (F (3,32) = 18.15, p <0.05). A posthoc comparison between the subgroups revealed that the number of carbon sources utilized by subgroup D was significantly higher than subgroup A, B, and C (Tukey HSD, p <0.05). All of the tested *Gardnerella* isolates were able to utilize pyruvic acid, palatinose, and L-rhamnose. The next most frequently utilized carbon sources were D-fructose (32/36, 97%) and L-fucose (32/36, 97%).

**Fig. 4.**
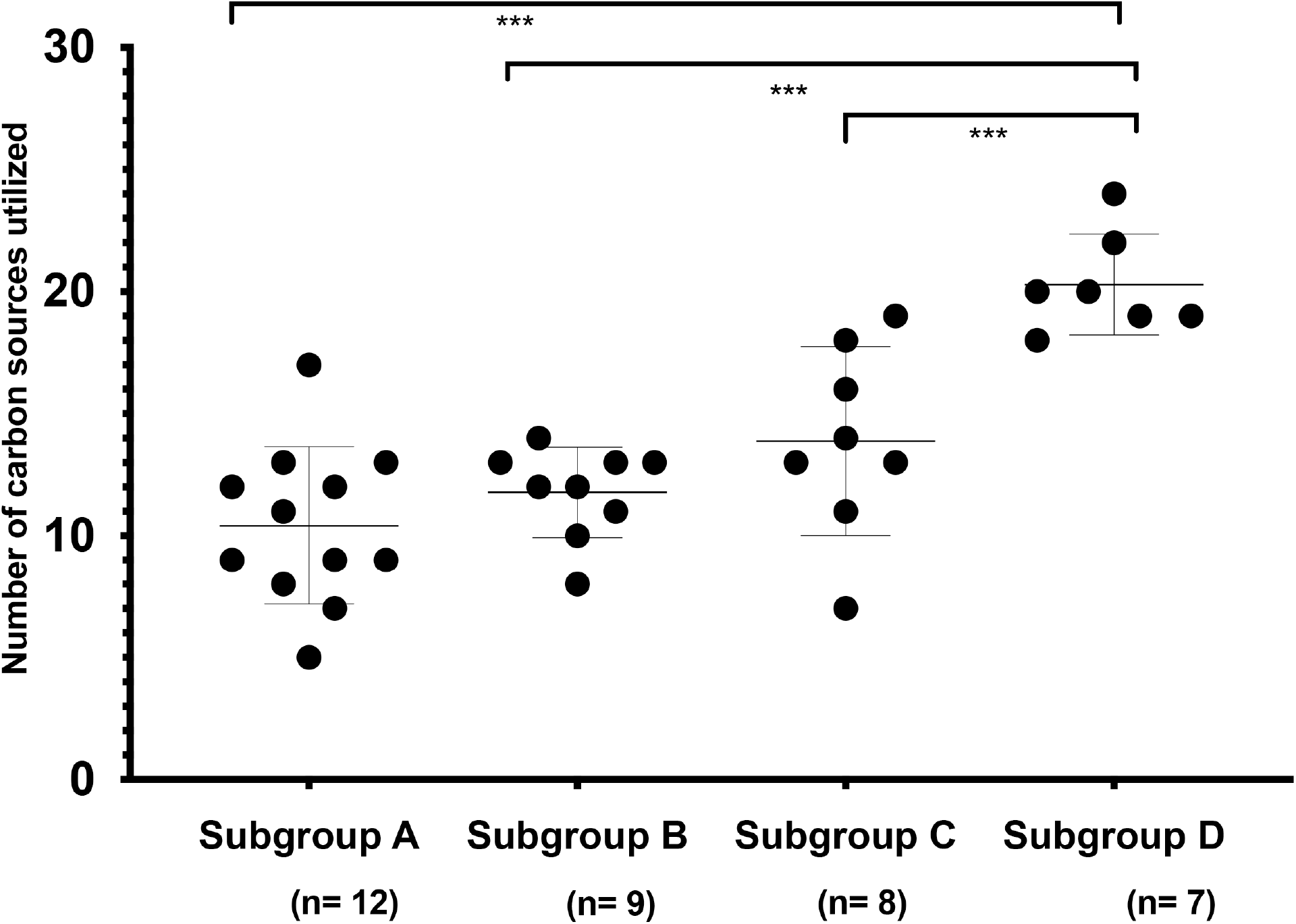
Comparison of numbers of carbon sources utilized by isolates in each subgroup. The number of carbon sources utilized by isolates in subgroup D was significantly higher than those in subgroups A, B and C (TukeyHSD, p ≤ 0.001). Only one fourth (9/36) of the tested isolates utilized more than 17 carbon sources, including all seven tested isolates of subgroup D.

Overall, 31/95 carbon sources were utilized by at least one isolate, and the majority (20/31) of these were sugars (mono-or oligosaccharides). Together, subgroup D isolates (n = 7) utilized more of the sugar substrates (18/37 available) than any other subgroup, including subgroup C (n=8), which utilized 15/37 available sugars. Utilization of any of the 11 available amino acids was rarely observed, with only two of the subgroup C isolates positive for L-methionine or L-valine utilization.

### Overlap in carbon sources utilization among the subgroups

To determine if subgroups could be distinguished based on carbon source utilization profiles, a principal component analysis was performed (Fig. 5). The overlap between the representative isolates of subgroups A, B, and C was significant. Subgroup D was significantly dissimilar to subgroups A and B (Fig. 5, pairwise-ADONIS, A vs D, R^2^ = 0.55; B vs D, R^2^ = 0.55, p<0.05). Although the dissimilarity between subgroup C and D was not statistically significant after Bonferroni adjustment, 39% (pairwise ADONIS, C vs D, R^2^= 0.39) of the variation in carbon source utilization could be explained by subgroup affiliation of the tested isolates, which was higher than between subgroups A, B and C (A vs B: 13%, A vs. C: 21%, and B vs. C: 8%).

**Fig. 5.**
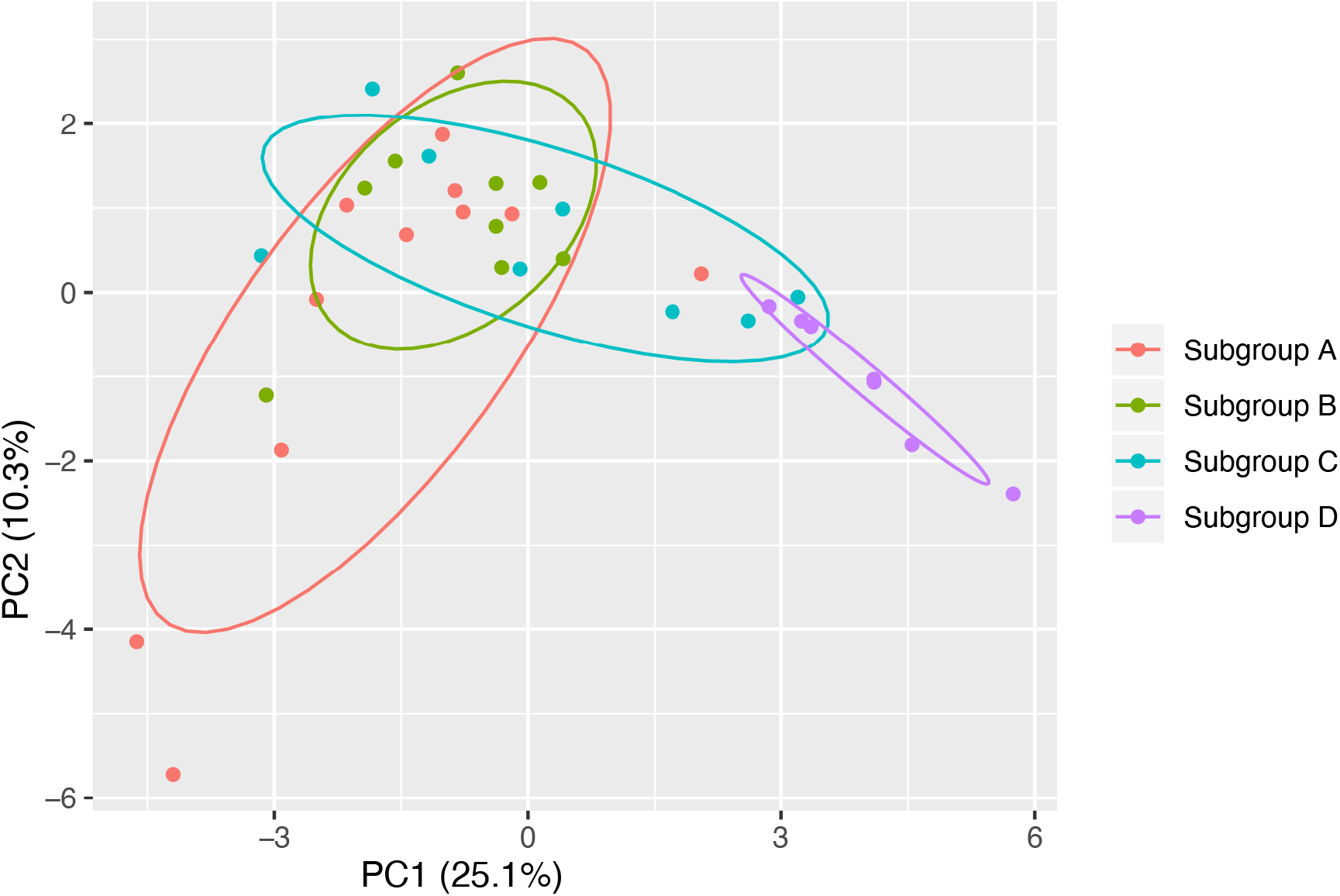
Subgroup D has minimal overlap with the other subgroups in carbon source utilization. The degree of variation based on carbon source utilization between subgroup D and subgroups A and B was significant (pairwise ADONIS, p < 0.05, after Bonferroni adjustment). The variation in carbon sources utilization between subgroup C and D can be explained by subgroup affiliation in 39% of cases. Overall, 42% of differences in carbon sources utilization between subgroups can be explained by their subgroup affiliation (Adonis, R^2^ = 0.42, p < 0.05).

### Association of carbon source utilization pattern with subgroups

To identify carbon sources that differentiate the subgroups, we selected twelve substrates that were utilized by more than five isolates but fewer than thirty isolates. Chi-square tests were performed to determine if the subgroups significantly differ in the utilization of those twelve carbon sources. The four *Gardnerella* subgroups differed in their use of 3 of the 12 carbon sources: turanose, inosine, and uridine 5-monophosphate (Chi-square test, p <0.05, Bonferroni adjusted) (Fig. 6). For each of these three carbon sources, subgroups A and B had low frequency of use (9.5% = 2/21, 0.0% = 0/21, 0.0% = 0/21; subgroups A and B combined), subgroup C had low or intermediate frequency of use (25.0% = 2/8, 50.0% = 4/8, and 62.5% = 5/8), whereas subgroup D had high frequency of use (100.0% = 7/7, 100.0% = 7/7, 100.0% = 7/7).

**Fig. 6.**
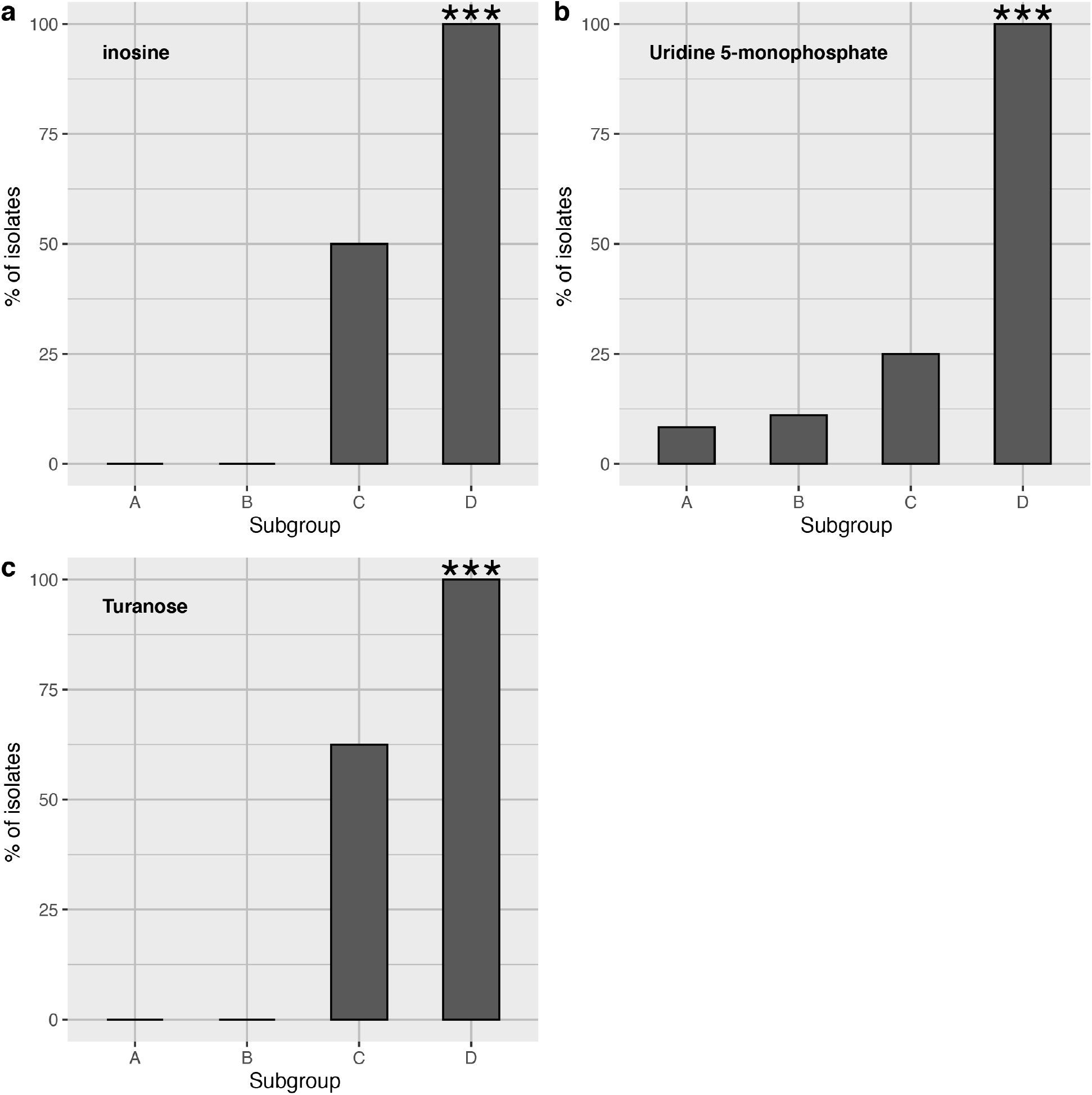
Subgroup D can be differentiated from the other subgroups based on its capacity to utilize inosine **(a)**, uridine 5-monophosphate **(b)**, and turanose **(c)**. The percentage of isolates in each subgroup that utilize the indicated carbon source is shown. The utilization of these three carbon sources is significantly associated with subgroup D (Chi-square test, p ≤ 0.001).

### Carbon source utilization by co-cultured isolates

Since the four subgroups co-exist in the same ecosystem, it is possible that mixing them might facilitate the utilization of certain carbon sources. To detect any such facilitation in carbon sources utilization, we co-cultured isolates from all four subgroups in six pairwise combinations (A+B, A+C, A+D, B+C, B+D, and C+D). The representative isolates of subgroups A-D utilized 11, 13, 19 and 24 carbon sources, respectively, when grown alone while co-cultures utilized from 12 to a maximum of 22 carbon sources (Table 1). In every case, the co-culture utilized fewer carbon sources than the isolate that utilized the most carbon sources on its own.

**Table 1.** Carbon source utilization by co-cultured isolates

| Isolate 1<br>(Subgroup) | Isolate 2<br>(Subgroup) | No. carbon sources |  |  |
| --- | --- | --- | --- | --- |
|  |  | Isolate 1 | Isolate 2 | Co-culture |
| VN003 (A) | VN002 (B) | 11 | 13 | 12 |
| VN003 (A) | NR001 (C) | 11 | 19 | 13 |
| VN003 (A) | WP012 (D) | 11 | 24 | 22 |
| VN002 (B) | NR001 (C) | 13 | 19 | 15 |
| VN002 (B) | WP012 (D) | 13 | 24 | 20 |
| NR001 (C) | WP012 (D) | 19 | 24 | 19 |

## Discussion

Rarely abundant species can be maintained in the human microbiome through a variety of mechanisms, which include but are not limited to sequestration of essential nutrients from competing species, diversification of phenotype [29], social cheating [30], and negative frequency dependent selection [17]. Differences in nutrient utilization among community members can be a key factor that sets the stage for negative frequency dependent selection [31]. The reproductive fitness of nutritional specialist species will remain high as long as the supply of nutrients usable by the specialists is abundant. As soon as the supply of these nutrients drops, slower-growing generalists, by virtue of their greater utilization capacity, will have increased fitness, which will eventually lead to their dominance in the absence of any other negative influences.

Among the four subgroups of *Gardnerella* spp. that colonize the vaginal microbiome of reproductive-aged women, subgroup D is the rarest in terms of abundance and prevalence among women [4]. Subgroup D is also relatively slow-growing, yet shows an increased growth rate when the number of competitors in an *in vitro* community increases [12]. We have reported previously that resource-based competition is common among *Gardnerella* spp. and no evidence for contact-dependent interaction was observed. Therefore, we set out to investigate if negative-frequency dependent selection is responsible for persistence of subgroup D, which would require relatively small niche overlap and a more generalist lifestyle than the other *Gardnerella* spp. in the vaginal microbiome.

### Predicted niche overlap between the four subgroups

Niche overlap may lead to competition for nutrients and space [31–34] and it has been reported that competition is prevalent among metabolically similar bacterial species [14]. Occupying distinct niches can therefore help bacterial species avoid competition for space, growth factors, and nutrients, resulting in increased reproductive fitness. Since subgroup D isolates have higher growth rates *in vitro* in the presence of competitors compared to when grown alone, these isolates presumably occupy a distinct niche. The pangenome analysis showed that the four subgroups differ significantly based on the composition of their predicted proteomes (Fig. 1), with only 176 proteins comprising the strict core of proteins found in all isolates. This finding is not surprising since the genetic diversity among *Gardnerella* is well established, and genome sequence comparisons formed the basis for the recent reclassification of *Gardnerella* into 13 genome species [1–3].

Comparisons of the entire predicted proteomes do not, however, focus on the key factors for a resource-based competition: nutrient utilization potential. Proteins involved in nutrient uptake and metabolism account for only a fraction of the 4,868 protein clusters comprising the pangenome. Analysis of the distribution of various functional (COG) categories of proteins revealed significant differences among subgroups in their predicted capacity to utilize carbohydrates and amino acids (Fig. 3a, 3b), with subgroup D having significantly more of its proteome dedicated to amino acid transport and metabolism than any of the other subgroups. Since a resource-based competition encapsulates competition for space, growth factors and nutrients, our findings from the pangenome and COG analyses suggest that the competition among the four subgroups is not spatial but may be primarily for nutrients; a speculation supported by the previous observation that *Gardnerella* spp. form multi-subgroup biofilms [12].

### Subgroup D is a nutritional generalist relative to subgroup A, B and C

The diversity of nutrients available to microbiota in the vaginal microbiome is less than in the gastrointestinal microbiome, where food intake provides a constant source of diverse nutrients that affect the assembly of gut microbiota [35, 36]. Vaginal microbiota, on the contrary, are largely dependent upon host-derived nutrients, the most abundant of which is glycogen. Glycogen is deposited in the vaginal lumen by epithelial cells under the influence of estrogen [37], and is digested into maltooligosaccharides, maltodextrins and glucose by the combined activities of host and microbial enzymes prior to uptake and metabolism by the microbiota [38–40]. Given the relatively narrow range of nutrients available in the vaginal microbiome, it is expected that the resident microbiota, including the four subgroups of *Gardnerella,* overlap to a considerable extent in their nutrient utilization capacity, resulting in some level of competition among them [32, 36, 41]. As discussed earlier, subgroup D is an exception since the growth of these isolates was actually facilitated in co-cultures, suggesting that while it may compete with other *Gardnerella* spp. over common nutrients like the breakdown products of glycogen, it may be able to utilize a greater overall diversity of nutrients (i.e. it is a generalist).

The AN microplate assay results showed that subgroup D isolates utilized more of the provided carbon sources than isolates in the three other subgroups (Fig. 4). Furthermore, when the patterns of substrate use were considered, subgroup A, B and C were not separable from each other, but subgroup D was significantly different (Fig. 5). The distinct pattern observed in subgroup D was partially driven by utilization of three particular substrates: turanose, inosine, and uridine 5-monophosphate (Fig. 6). Turanose is an isomer of sucrose, known as a non-accumulative osmoprotectant, aiding bacterial growth at high osmolarity [42]. The importance of turanose utilization in the vaginal environment is not known yet, but our observation is an indication that subgroup D isolates can metabolize sucrose-like sugars. The two other carbon sources: inosine and uridine 5-monophosphate are probably used in purine and pyrimidine biosynthesis in *Gardnerella* spp‥

Some findings of the pangenome and COG analyses could not be reconciled with the phenotypic carbon source utilization assay. For example, although subgroup C and D have higher proportions of their proteomes predicted to be involved in transport and metabolism of carbohydrates (Category G) and amino acids (Category E), respectively, than the other subgroups, subgroup C isolates did not utilize the greatest number of available sugar substrates in the AN microplate and subgroup D isolates did not utilize any of the amino acid substrates available. It is, however, important to consider that the carbon source utilization assay was performed in a plastic environment and included only 95 substrates, many of which are not relevant to the vaginal microbiome. More relevant amino acid sources available in the vagina, including those whose abundance is altered in bacterial vaginosis, such as isoleucine, leucine, proline, and tryptophan, are not included [43–45]. Ideally, this study would have involved a vaginal-microbiome specific nutrient panel, but such reagents were not available. Even with this limitation, our results suggest that subgroup D is a nutritional generalist relative to other *Gardnerella* spp‥ Most of the ecological studies that have been performed to date to elucidate mechanisms shaping the assembly of bacterial communities have included either environmental bacterial species or well-characterized model organisms [14, 16, 29, 36, 41, 46–50]. There are understandably fewer studies that focus on interactions among host-associated microbiota [51, 52].

### Negative-frequency-dependent selection in the vaginal microbiome

The genomic and phenotypic differences we observed between subgroup D and the three other subgroups, including the potential to utilize more amino acids, use of a greater number of carbon sources, and a distinct pattern of substrate utilization, suggest that subgroup D is a candidate for negative frequency dependent selection. Why then are these *Gardnerella* spp. only observed rarely, and in low abundance in reproductive aged women? Among 413 vaginal samples from reproductive aged Canadian women, genome species comprising subgroup D of *Gardnerella* were detected in <10% of samples and never accounted for more than 5% of the microbiota [4]. Vaginal environmental dynamics and related host factors, such as menstruation, sloughing of epithelial cells, and fluctuating pH contribute to the turnover of bacterial species, shifting the bacterial population density and changing the nutrients available [53]. A decline in population density would reshuffle the vaginal ecosystem, increasing the supply of abundant nutrients accessible to faster growing, specialists, and checking the growth of slower growing generalist subgroup D.

Although subgroup D is likely rare due the factors described above, it could still be a major player in ecological succession and transition of vaginal microbiota between a *Lactobacillus* dominated community and the overgrowth of anaerobes characteristic of bacterial vaginosis. These organisms may also play a particular role in biofilm formation or competition for occupancy of the vaginal mucosa. Rarely abundant species often act as keystone species helping colonization by other bacterial species, which are also essential to maintain homeostasis of an ecosystem [54–57]. Resolution of the role of low abundant *Gardnerella* spp. will depend on the development and application of experimental systems that more closely model the human vaginal microbiome. Rodent models have shown some promise, especially for studies of specific combinations of organisms [58], but there is also potential in bioreactors [59], and cell and tissue culture systems that attempt to recapitulate many of the environmental and physiological aspects of the vaginal microbiome [60]. Further study of rare *Gardnerella* spp. will likely also result in the definition of additional species within this diverse genus.

## Acknowledgements

The authors are grateful to Champika Fernando for excellent technical support and to all members of the Hill Lab for their review of the manuscript. Special thanks to Prairie Diagnostic Services for access to their anaerobic chamber. No thanks to COVID-19.

## Declarations

### Funding

The research was supported by a Natural Sciences and Engineering Research Council of Canada Discovery Grant to JEH.

### Competing interests

None declared.

### Availability of data and material

NCBI Bioproject accession numbers for all genome sequence data are included in Table S1.

### Authors’ contributions

Conceived and designed the study: Salahuddin Khan and Janet E. Hill. Performed the experiments: Salahuddin Khan and Sarah J. Vancuren. Analysed the data: Salahuddin Khan, Sarah J. Vancuren, Janet E. Hill. Wrote and revised the manuscript: Salahuddin Khan, Sarah J. Vancuren, Janet E. Hill.

---

**Fig S1.**
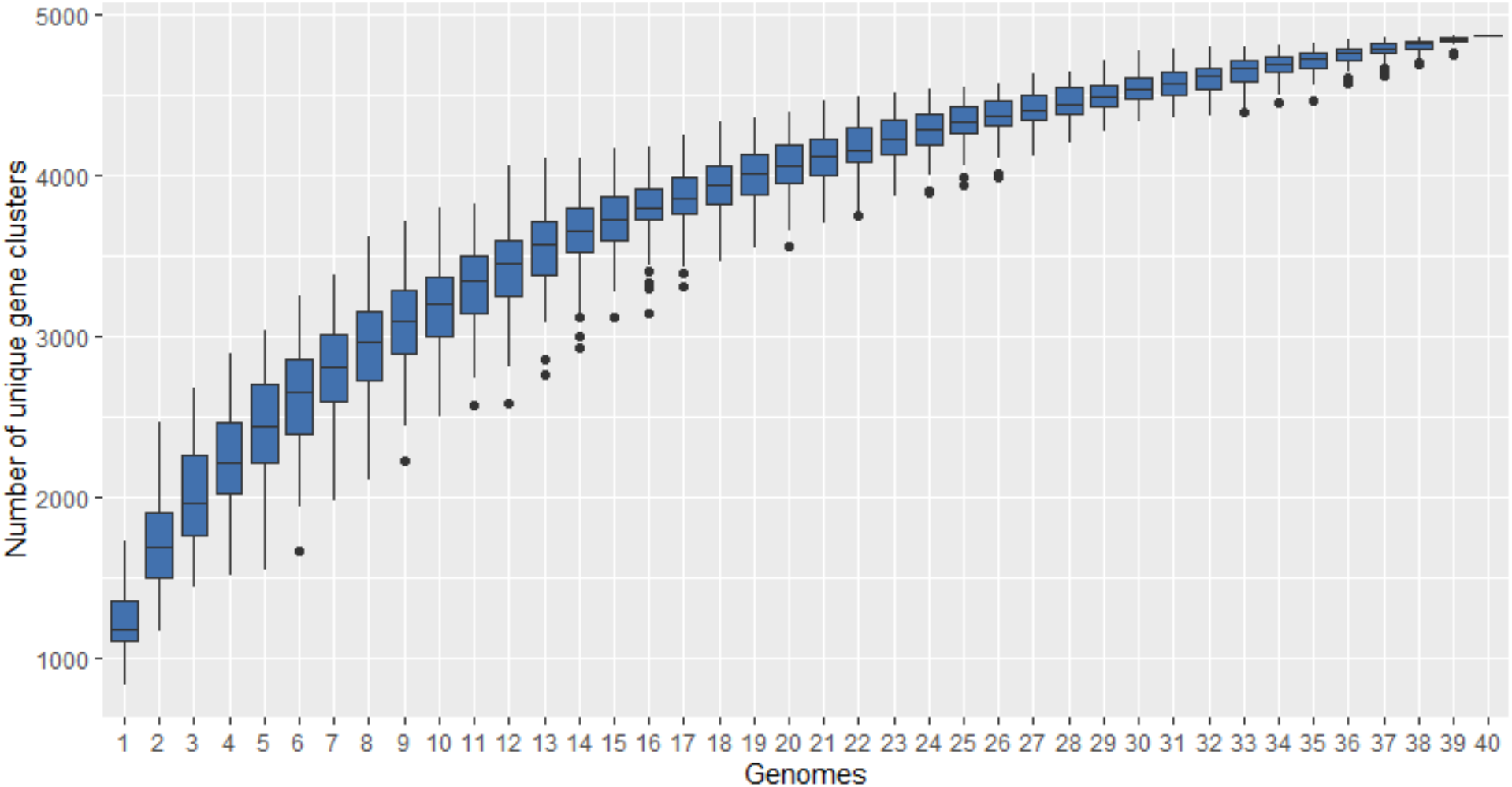
Rarefaction curve of the pangenome of *Gardnerella* spp‥

**Table S1.**
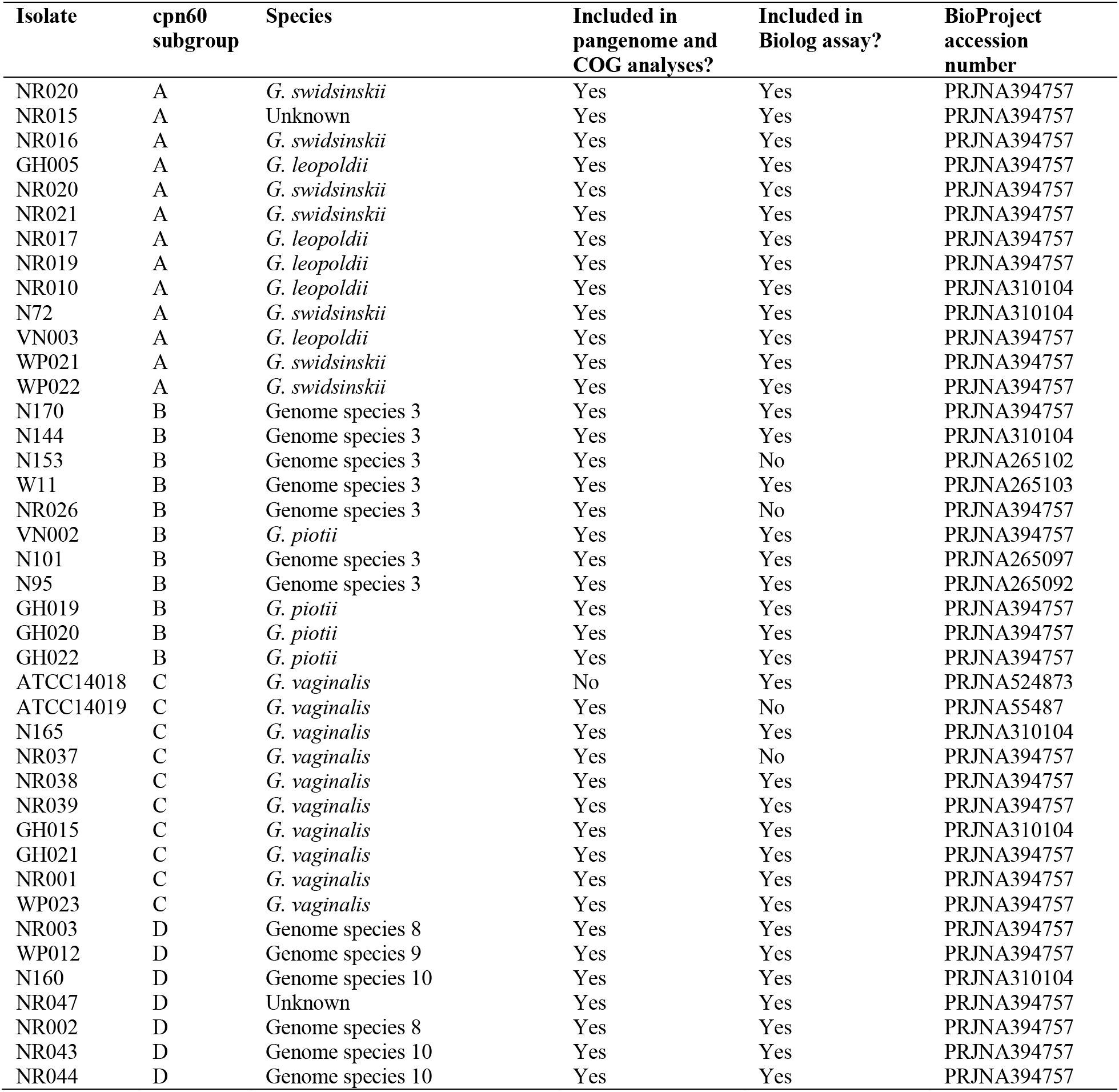
Bacterial isolates and whole-genome sequences used for pangenome analysis and carbon source utilization profiling assay.

## Notes

### Competing Interest Statement

The authors have declared no competing interest.

### Summary of Updates

Figures 1, 2, 3 and 6 revised. Table 1 added.

